# Lipopolysaccharide-induced neuroinflammation disrupts functional connectivity and community structure in primary cortical microtissues

**DOI:** 10.1101/2021.07.08.451705

**Authors:** Elaina Atherton, Sophie Brown, Emily Papiez, Maria I. Restrepo, David A. Borton

## Abstract

Three-dimensional (3D) neural microtissues are a powerful in vitro paradigm for studying brain development and disease under controlled conditions, while maintaining many key attributes of the in vivo environment. Here, we used primary cortical microtissues to study the effects of neuroinflammation on neural microcircuits. We demonstrated the use of a genetically encoded calcium indicator combined with a novel live-imaging platform to record spontaneous calcium transients in microtissues from day 14-34 in vitro. We implemented graph theory analysis of calcium activity to characterize underlying functional connectivity and community structure of microcircuits, which are capable of capturing subtle changes in network dynamics during early diseases states. We found that microtissues cultured for 34 days displayed functional remodeling of microcircuits and that community structure strengthened over time. Lipopolysaccharide, a neuroinflammatory agent, significantly increased functional connectivity and disrupted community structure 5-9 days after exposure. These microcircuit-level changes have broad implications for the role of neuroinflammation in functional dysregulation of neural networks.

## Introduction

Three-dimensional (3D) cell culture techniques have been developed to create brain-like microenvironments to study neural development and disease at the cellular level. Compared to traditional two-dimensional (2D) monolayer cultures, 3D cultures have been shown to more closely recapitulate features of the in vivo environment, including morphology, cell-cell interactions, and cell signaling^1–3^. Further, 3D neural cultures are capable of forming complex structural and functional networks, which is vital to understanding neural behavior and cannot be recapitulated in two dimensions (2D)^4–6^.

3D culturing strategies vary widely in both scaffolding and cell source. The use of culturing scaffolds can create a well-controlled structural framework for seeded cells ^7–10^, some of which have been shown to facilitate guided axon growth^11^ and even capillary-like delivery of nutrients^12^. Alternatively, scaffold free cultures produce an endogenous extracellular matrix, which impacts neural network formation, cell signaling, migration and mimics the mechanical properties of brain tissue^1,2,13–15^. Normal brain function, as well dysfunction, is resultant from a complex interplay between many different cell types, and as such, in vitro studies have moved toward multicellular models. These models can be achieved through a number of culturing strategies, including co-culturing of immortalized cell lines, heterogeneous stem cell differentiation, or primary cell collection from a multicellular tissue source^16–18^. Leveraging a primary cell collection strategy, we created self-assembled 3D neural cultures sourced from rat cortical tissue, which incorporate many endogenous cells types and develop spontaneous electrical activity in neurons^13^.

Formation of coordinated neural network activity is a primary function of a healthy brain, and thus, is an important feature to characterize in an in vitro model of the brain. Recording longitudinal neural activity in vitro has predominantly been achieved with microelectrode arrays (MEAs)^19^. MEA’s are capable of capturing individual action potentials from cells cultured on top of flat electrode pads^20^. MEA platforms have been used in vitro to show large developmental changes in neural activity over weeks^15,19–23^. While MEA’s provide high temporal resolution of neuronal activity, genetically encoded calcium indicators (GECI) provide superior spatial resolution of functional data^24,25^. Calcium imaging is particularly well-suited to the in vitro environment because cultured neurons are not optically obstructed by the meninges or skull as they are in vivo. Calcium imaging has been used in vitro to examine functional neural behavior under controlled conditions to study both development and disease^3,7,23,26–30^.

As neural behavior emerges out of the formation of diverse hierarchical structural and functional networks, evaluating network-wide activity is essential to understanding in vitro microenvironments. Graph theory analysis (GTA) is a mathematical method of characterizing complex network interactions and has been used to examine the underlying FC (FC) and community structure of neural networks in both the in vivo and in vitro setting^31^. These “neuronal graphs” have primarily been applied to neural “mesocircuits”, in which nodes are spread across multiple brain areas in order to characterize regional brain connectivity from fMRI and electrical recordings in vivo^31^. At the scale of single cells, GTA of calcium transients provides quantified measures of connectivity within neural “microcircuits”^32^. Changes at the scale of microcircuits may be important in understanding early disease states, as functional changes of microcircuitry can be detected prior to onset of primary disease associated behavioral deficits^27,28,33^.

Here, we demonstrate a multimodal approach to collecting and characterizing longitudinal microcircuit activity in a 3D cortical primary culture for the application of studying neural development and disease in vitro. We dissociated primary rat cortical cells and seeded them into a custom injection-molded non-adherent agarose microwell, which leverages previously developed 3D self-assembled culturing methods^13,34^. AAV mediated transgene delivery of the GCaMP6s calcium indicator^24^ facilitated visualization of spontaneous fluorescent calcium transients that emerged by day in vitro (DIV) 14. We demonstrate the use of this culturing method for studying neural microcircuit development from day 14 to 34 in vitro. GTA of longitudinally imaged microtissues exposed the formation of functionally connected networks of neurons in the 3D primary culture. Furthermore, GTA revealed functional remodeling that occurred between week 2 (DIV14-20), 3 (DIV21-27), and 4 (DIV28-34) in vitro. Network changes were accompanied by an increase in modularity, an indication of community structure maturation described by the formation of highly connected groups of cells called “modules”. We found that correlation of single-cell node-pairs within modules increased, while correlation of nodes pairs between modules decreased, resulting in the strengthening of community structure. We then aimed to study the effect of acute neuroinflammation on dynamic functional properties of the microtissue networks by introducing lipopolysaccharide (LPS) to the cultures. LPS is a bacterial endotoxin known to induce neuroinflammation via changes to glial morphology, secretome, and neuronal fate^18,35–37^. We found no change in microcircuit FC from 2 hours to 3 days following LPS exposure, but a significant increase in FC at 5-9 days post exposure. This increase in FC disproportionately increased the inter-modular correlations, reducing modularity and the strength of community structure. Leveraging GECI for longitudinal functional imaging, we used graph theoretical analysis to reveal subtle but significant changes during development and the progression of LPS-induced neuroinflammation.

## Results

### Culturing platform for longitudinal live imaging

For our in vitro experiments, we utilized a three-dimensional, self-assembled, primary cortical culture previously shown to contain myriad cell types and produce an endogenous extracellular matrix^13,14^. To facilitate live imaging, we adapted a spheroid cortical culture model into a “trampoline” shaped microtissue, previously described by Schell et. al^34^. The trampoline shape, which maintains a heterogeneous cell composition (Supplemental Figure 1), provides a larger surface area compared to spheroids, increasing the number of imageable cells within the z-axis constraints of the microscope.

As high magnification objectives have very limited optical working distances, 3D cultures typically require physical manipulation to bring them closer to the objective by either transferring into a glass bottom imaging dish or removal of media to prevent samples from floating. As neural cultures are sensitive to environmental changes^20^, we aimed to collect image data without disrupting the culturing environment. To achieve this, we designed a custom agarose mold to place the cells within 350um of the bottom of the plate and, importantly, within the working distance of high-power objectives. Our mold is built upon published work utilizing agarose “stamping” for live imaging of spheroids^38^. We created a custom machined stainless steel injection mold consisting of a hollow tube with an endcap containing 6 machined holes to form the agarose pegs (Figure 1A). The injection mold is placed into a single well of a multi-well culture plate, suspended by a 3D printed holder (Figure 1B). Hot liquid agarose can be pushed through the injection mold and into the well with a syringe (Figure 1C). When the agarose cools, the injection mold can be removed (Figure 1D), leaving behind the agarose pegs within the microwell (Figure 1E). After primary tissue collection and dissociation, cells were seeded into the microwell (Supplemental Figure 3), placing them approximately 350um from the bottom of the plate and within the working distance of the objective for live imaging (Figure 1D).

**Figure 1.**
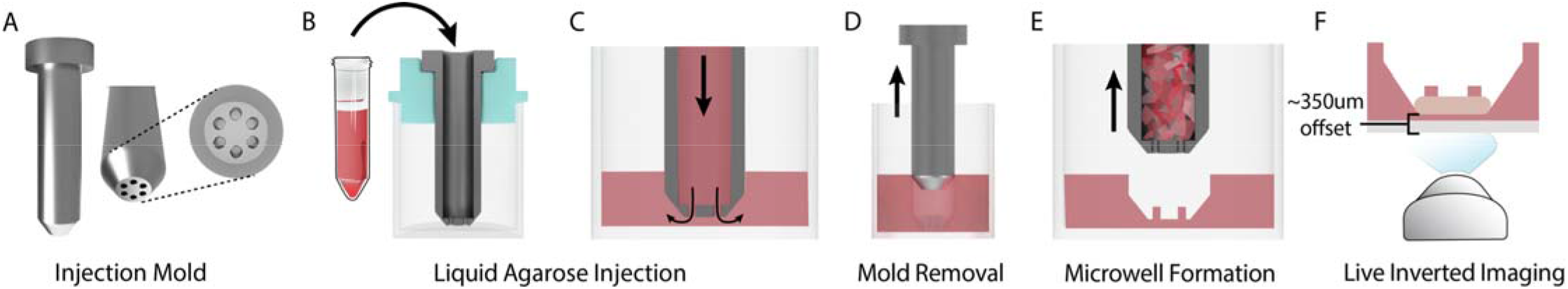
Custom injection mold positions the microtissue for live inverted imaging without the need for physical manipulation of tissues. (A) The injection mold design consists of a hollow tube with patterned end cap. The 6 holes in the endcap will form the 600um diameter agarose pegs of the microwell. (B) A cross section image of the injection molding apparatus. The injection mold (gray) is placed in the well, suspended to the correct height by a 3D printed holder (turquoise), and hot liquid agarose (red) is pushed into the hollow tube of the injection mold with a sterile syringe. (C) Liquid agarose moves through holes in the endcap and into the well. (D) When the agarose (red) is cooled, the mold can be gently pulled upward and removed. (E) Removal of the injection mold reveals the microwell with 6 pegs to produce the “trampoline” tissue shape. (D) This process places the microtissue (tan) around 350um from the bottom of the glass plate, within the working distance of most objectives.

### AAV mediated expression of GCaMP6s reveals spontaneous neural activity

Once microtissues were constructed, we applied a commercially available GECI to visualize neural activity in vitro. The GCaMP6s calcium indicator was expressed through AAV mediated transgene delivery (AAV1-hSyn1-mRuby2-GSG-P2A-GCaMP6s-WPRE-pA, AddGene). Expression of the fluorescent protein packaged in the viral capsid was driven under the human synapsin (hSyn1) promoter, as it produces neuron-specific expression patterns^39^. The GCaMP6s probe was chosen over GCaMP6f due to its higher signal to noise ratio^24^, which we expected to compensate for any optical interference from agarose or tissue itself.

After culturing microtissues for 1 day (Figure 2A), fluorescence expression was achieved via transgene delivery by culturing in virus-containing media for 3 days (Figure 2B). Initial fluorescence appeared at DIV 8 and progressively increased in brightness through DIV 14 (Figure 2C). Onset of spontaneous calcium transients were observed and recorded starting at DIV 14 (Figure 2D). Python code was used to identify cell bodies by applying Laplacian of Gaussian filtering to maximum projections of time-lapse images and non-maxima suppression to detect individual regions of interest (ROIs, Figure 2E). ROIs were curated to remove cells outside of the microtissue edge using manual interactive correction. Using curated ROI seed positions, ROI masks were created and overlaid onto each image of the time-lapse videos (Figure 2F) in MATLAB, and calcium transients and events were extracted from the masked videos (Figure 2G) with the FluoroSNNAP^40^ application.

**Figure 2.**
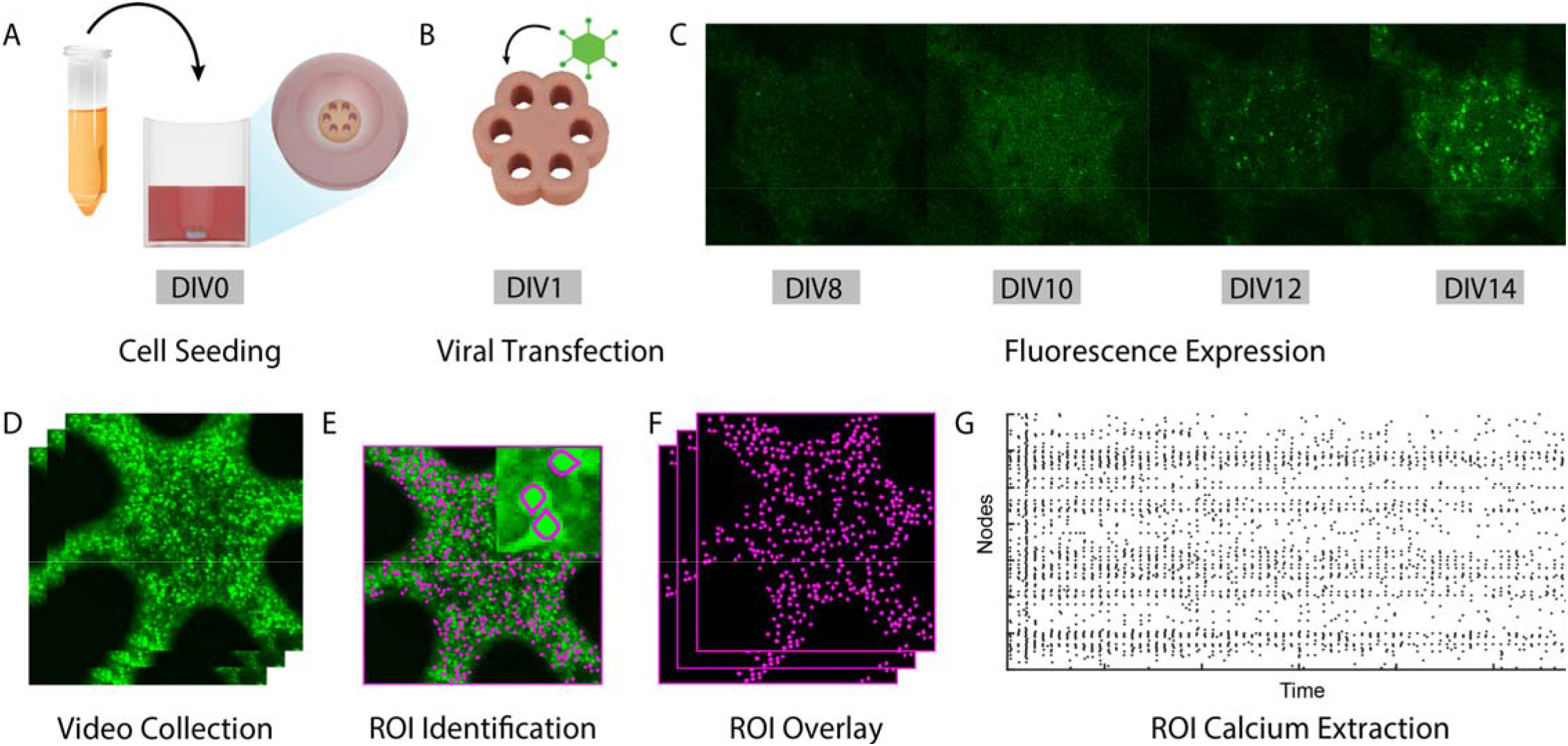
Three-dimensional self-assembled neural cultures express stable GcaMP6s fluorescence and exhibit spontaneous activity by DIV 14. (A) Dissociated P1 primary rat cells are seeded into a custom injection molded agarose well. (B) Cells, which form a “microtissue” (tan) in the first 24 hours, were transfected with a commercially available viral vector to encode the GcaMP6s calcium indicator (AAV1-hSyn1-mRuby2-GSG-P2A-GcaMP6s-WPRE-pA). (C) Microtissues display progressive expression of baseline calcium fluorescence from DIV 8 – DIV14. (D) Time-lapse videos of calcium transients were collected for 4 minutes. (E) Python code was used to perform semi-automated segmentation of cell bodies. (F) Segmented regions of interest (ROIs) were overlaid onto the 4-minute video for single-cell calcium trace extraction. (G) A raster plot of calcium events detected from calcium traces were extracted from segmented videos using the FluoroSNNAP application^40^.

### Microtissues exhibit significant remodeling of FC during development

We characterized FC of microtissues from onset of spontaneous calcium transients. To accomplish this, we recorded activity in 9 microtissues across three biological replicate litters from DIV14-34. We analyzed network activity from recordings using graph theory analysis (GTA), a mathematical method of describing complex interactions between “nodes’’ in an interconnected network^31^. Here, we employed GTA to describe in vitro network activity at the microcircuit level, where each node within the graph represents a single cell^32,41^. Using a network analysis toolkit^7^, we characterized correlation coefficients, clustering coefficients, and path lengths of microcircuits. To assess trends in neural activity over time, recordings were grouped into weeks (week2=DIV14-20, week3=DIV21-27, week4=DIV28-34), where each data point represents a single recording of a microtissue within the time range.

The correlation coefficient is a metric of FC between pairs of nodes. Correlation was determined by the statistical similarity of activity between each node pair by Pearson cross-correlation. Higher correlation coefficients indicate stronger functional connections between nodes^32^. We assessed the average correlation of microcircuits, calculated by taking the mean correlation coefficient of all node-pairs. We found a significant (p=0.0026) reduction in the average correlation from week 2-3 and a significant (p=0.0151) recovery in week 4 (Figure 3A).

**Figure 3.**
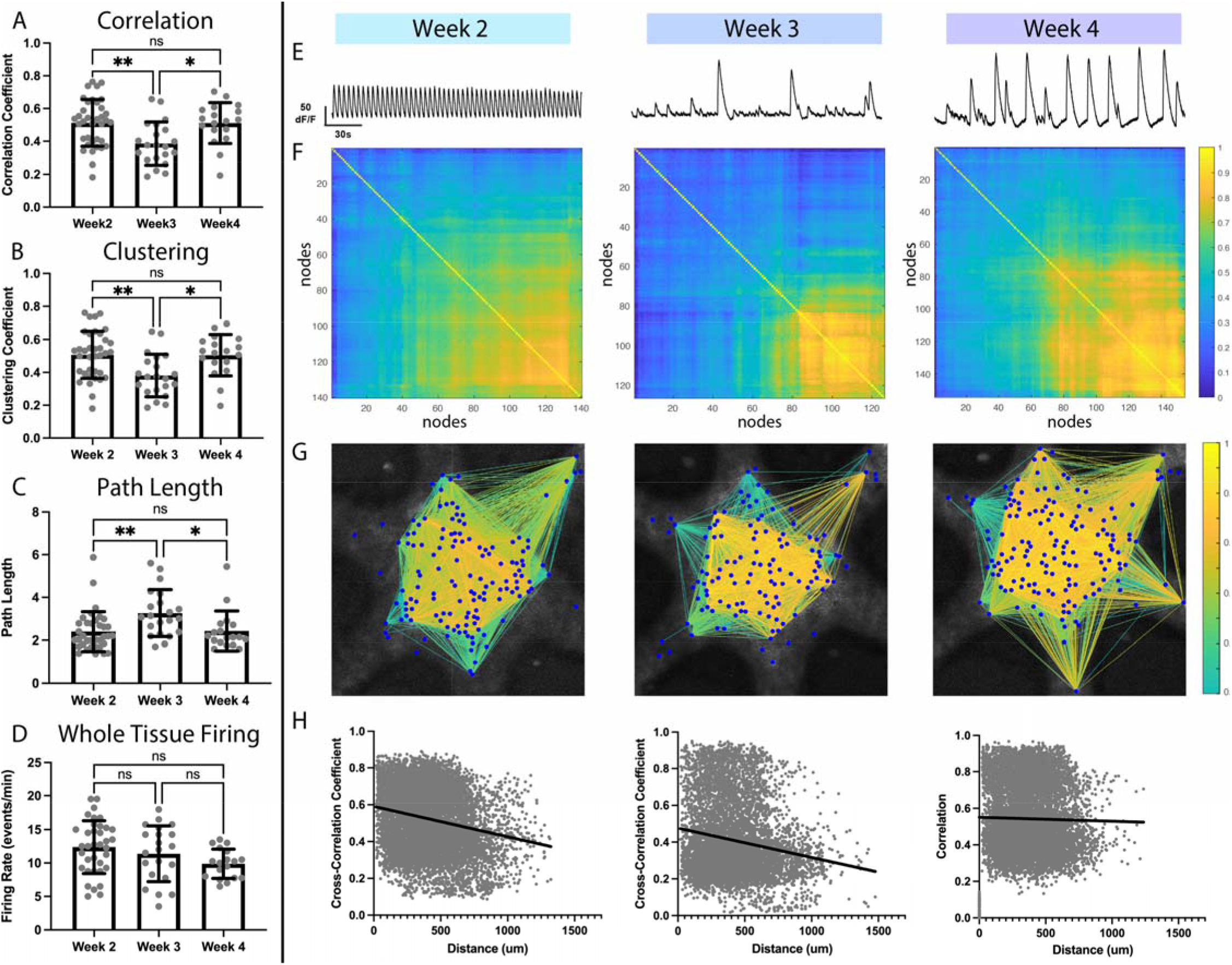
Spontaneous calcium activity reveals significant functional remodeling over three weeks of development in vitro. (A) Average correlation coefficients of microtissues significantly decrease (p=0.0026) from week 2 to week 3, followed by a significant increase (p=0.0151) from week 3 to week 4. (B) Clustering coefficients follow the same trend as the correlation, with a significant decrease (p=0.0032) of average clustering coefficient in week 3 and a subsequent significant increase (p=0.016) in week 4. (C) Path length between nodes followed the inverse trend, with a significant increase (p=0.0054) in average path length in week 3 and a significant decrease (p=0.0263) in week 4. (D) Whole-tissue firing rate showed no significant differences (week2-week3, p=0.5851; week 3-week 4, p=0.4209) over 3 weeks of development. (E) Whole-tissue calcium traces recorded in a single example microtissue from week 2, week 3, and week 4 show changes in firing patterns over time. (F) Correlograms from pairwise Pearson cross-correlations coefficients between nodes from recordings shown in (E), exhibit progression of node connectivity across weeks. (G) Correlational connectomes overlay the cross-correlation values above 0.5 onto the physical node positions. Line color connecting nodes correlates to the correlation coefficient value. (H) Plots of the correlation coefficients versus the physical distance between nodes, shows no preference for strong local connections over cross-tissue connections. Significance to compare multiple weeks was determined with a one-way ANOVA with p<0.05 (^*^p<0.05, ^**^p<0.01).

The clustering coefficient is a measure of FC between triplets of nodes, representing the formation of “small world” network organization^42^. A higher clustering coefficient corresponds to the presence of highly interconnected clusters of nodes, an indication of higher network complexity compared to a random network^31,43^. Average clustering coefficients followed the same trend as average correlations over 3 weeks in vitro, with a significant reduction (p=0.0032) from week 2-3 and a significant (p=0.016) recovery in week 4 (Figure 3B).

Path length is a metric of network integration, inversely related to the correlation and representing the ability of nodes to communicate efficiently with limited intermediary nodes^31,44^. Node-pairs with short path lengths are thought to be efficient at sharing information. We found a significant increase (p=0.0054) in the average path lengths from week 2-3 and a significant (p=0.0263) recovery in week 4, inversely following trends of the correlation and clustering (Figure 3C). Collectively, the disruption and recovery of correlation, clustering, and path length indicate significant remodeling of FC in microtissues during maturation.

Additionally, we investigated if these changes in FC were associated with changes in synchronous neuronal bursting. Thus, we examined whole-tissue firing rates, measured by the number of tissue-wide synchronous bursts per minute identified by the FluoroSNNAP event detector. While there was a decrease in the average whole-tissue firing rate, the changes were not statistically significant (week2-3, p=0.5851; week3-4, p=0.4209; Figure 3D). This finding suggests that global firing activity is not sensitive enough to capture the subtle synaptic changes of functional remodeling at this stage of maturation.

To better understand the changes in connectivity, we examined the longitudinal FC of individual microtissues. We found that individual microtissues displayed changes in firing patterns over time (Figure 3E, Supplemental Figure 3). Correlograms of a representative microtissue from each week (Figure 3F) showed that the net reduction in average correlations in week 3 did not reflect a blanket reduction in all node-pairs, marked by a visible increase in correlation for a subset of node-pairs. Correlational connectomes (Figure 3G), which overlay correlation values above 0.5 onto the physical microtissue, showed this increase in strongly correlated node-pairs (correlation>0.8) more clearly. This qualitatively supports the idea that microtissues undergo selective pruning of FC during maturation. To investigate the structural composition of the network, we then characterized the correlation of proximal nodes. We found that physical proximity of nodes was not a strong predictor of correlation in individual microtissues (DIV14, R^2^=0.45; DIV24, R^2^=0.030; DIV32, R^2^=0.00045, Figure 3H) indicating that the underlying structural network is likely not a regular network, which is primarily made up of proximal connections. These data further support the presence of significant whole-tissue remodeling over the first 4 weeks in vitro.

### Microtissues develop and refine community structure over time

We assessed the strength of microcircuit community structure, characterized by the formation of strongly connected groups of neurons within the functional network. The development of community structure is an important feature of neural network maturation both in vitro and in vivo.^19,45,46^ Here, we used a compiled network analysis toolkit^7^ to characterize community structure.

Modularity is a metric of community structure maturation defined by the presence of highly connected groups of nodes called “modules”^31,32,41^. Modules can be identified by hierarchical clustering of node-pair correlations^7,47^. As modularity increases, network segregation increases, resulting in strong node-pair connections within modules and weak node-pair connections between modules^47^. Modularity significantly increased (p=0.046) from week 2-3 (Figure 4A), despite the decrease in overall correlation, suggesting that functional remodeling coincides with construction of modules. Additionally, microtissue modularity remained high at week 4 (p=0.0475; Figure 4A), indicating that the increased correlation did not disrupt the community structure developed in week 3. Remarkably, modules did not represent physical “regions” within the microtissue. Nodes of different modules and intra-modular connections were interspersed (red, module1; blue, module2; Figure 4B).

**Figure 4.**
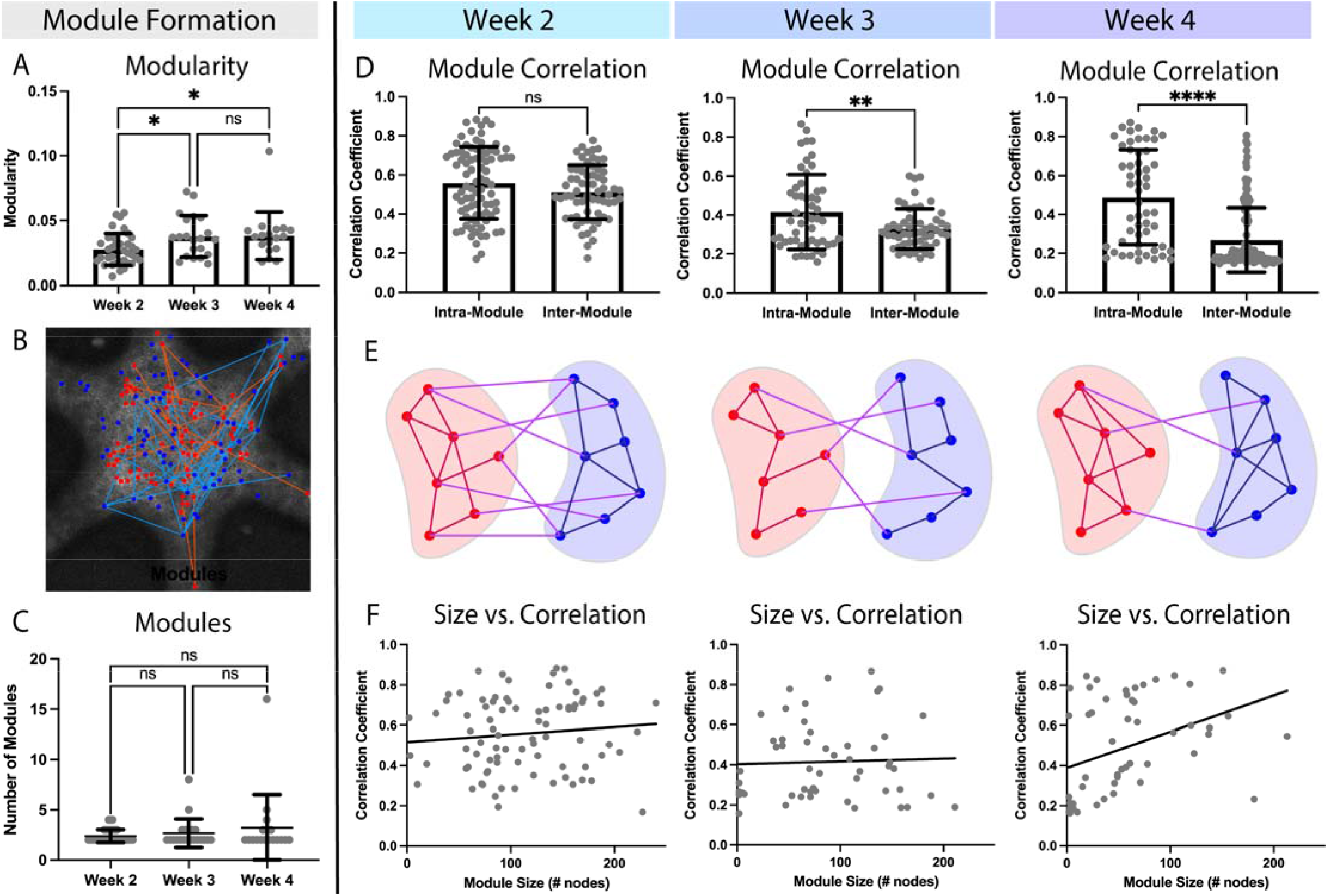
Microtissues develop community structure through selective functional remodeling. (A) Overall modularity of microtissues significantly increased (p=0.046) at week 3 and remained increased (p=0.0475) at week 4. (B) Representative image of the physical layout of module nodes. Module 1 nodes (red dots) and their intra-modular connections (orange lines) are physically interspersed with module 2 nodes (blue dots) and their intra-modular connections (light blue lines). (C) Number of modules present in microtissues does not change over the first 4 weeks of development. (D) The intra-module correlation is not significantly different (p = 0.0834) from the inter-module correlation in week 2. In weeks 3 and 4 the intra- and inter-module correlations significantly separated (week 3, p =0.0045; week 4, p <0.0001). (E) Schematics of community structure formation based on data from (D), with intra-modular connections in red (module 1) or blue (module 2), and inter-module connections in magenta. Week 2 reflects that there was no difference between the intra-module and inter-module connections. The week 3 schematic depicts the overall decrease in functional connectivity, which disproportionately affects inter-module connection. The week 4 schematic shows the overall increase in connectivity, with an increase in intra-module connections and a decrease in inter-module connections. (F) There is little correlation between module size (number of nodes) and the intra-module correlation at week 2 and week 3. At week 4 there is a small positive correlation between module size and average correlation, indicating that large modules in week 4 were functionally more connected. Significance for modularity and number of modules across multiple weeks were determined with a one-way ANOVA with p<0.05 (*p<0.05). Significance between intra-modular and inter-modular correlations were determined with unpaired, two tailed t-tests with p<0.05 (^**^p<0.01, ^***^p<0.001, ^****^p<0.0001).

Although the modularity changed over time, the absolute number of identified modules did not (week2-3, p=0.8449; week3-4, p=0.6127; Figure 4C), suggesting that increased modularity may be related to the segregation of established modules rather than further subdivision of modules. To investigate, we compared node-pair correlations within modules (intra-modular correlations) to those between modules (inter-modular correlations). Intra-modular and inter-modular correlations were not significantly different in week 2 (p=0.0834), but progressively separated in week 3 (p=0.0045) and 4 (p<0.0001; Figure 4D). These data indicating that microtissues at week 2 contained weak community structure in which connectivity within modules was equivalent to those between modules (Figure 4E). At week 3, the overall FC decrease disproportionately reduced the strength of inter-modular connections, resulting a separation of intra- and inter-modular correlations (Figure 4E). Further, the overall increase in connectivity in week 4 disproportionately increased intra-modular connections (Figure 4E), resulting in the emergence of well-defined community structure. Additionally, the number of nodes within strongly correlated modules increased over time (Figure 4F), with module size showing limited correlation to module connectivity in week 2 (R^2^=0.01134; slope deviation from zero, p=0.3320) and week 3 (R^2^=0.0015, slope deviation from zero, p=0.7818), but a stronger correlation in week 4 (R^2^=0.1462; slope deviation from zero, p=0.0043; Figure 4F). Collectively these data indicate a progressive strengthening of community structure created through selective pruning of FC over weeks in vitro.

### Functional connectivity significantly increases 5-9 days after LPS exposure

We then investigated FC of 3D in vitro neural networks during acute neuroinflammation. We applied the methods of functional imaging and analysis described above to a well-established lipopolysaccharide (LPS) model of acute neuroinflammation. LPS stimulates glial inflammatory cascades, producing changes in cellular morphology^18^, gene expression^48^, and secretome^35^. While LPS-induced neuroinflammation does not model a specific disease pathology, it can add to our understanding of basic neuroinflammatory cellular dynamics. LPS-induced neuroinflammation has significant downstream effects on neural function, including connectivity changes in mesocircuits^49–51^, the development of epileptiform hyperactivity^52^, and cognitive deficits in rodents and humans^49,53^. Here, we examined the effect of LPS on microcircuit FC in microtissues.

As LPS exposure in vivo has effects on the secretome in tissues directly after exposure^35^, morphology after 2 days^18^, and synaptic proteins at one week^54^, we examined FC at three corresponding timeframes (2hrs, 1-3 days, and 5-9 days) after exposure. Microtissues were treated with either 10ug/mL of LPS or a PBS control at DIV 25 and imaged at 2hrs after treatment (LPS = 9 microtissues; PBS = 8 microtissues, Figure 5A). Samples were subsequently imaged at 1-3 days (DIV26/27/28) and 5-9 days (DIV30/32/34). Due to the sensitivity of neuronal activity to any environmental changes, microtissues were allowed to recover from media changes for 2 hours prior to imaging^20^.

**Figure 5.**
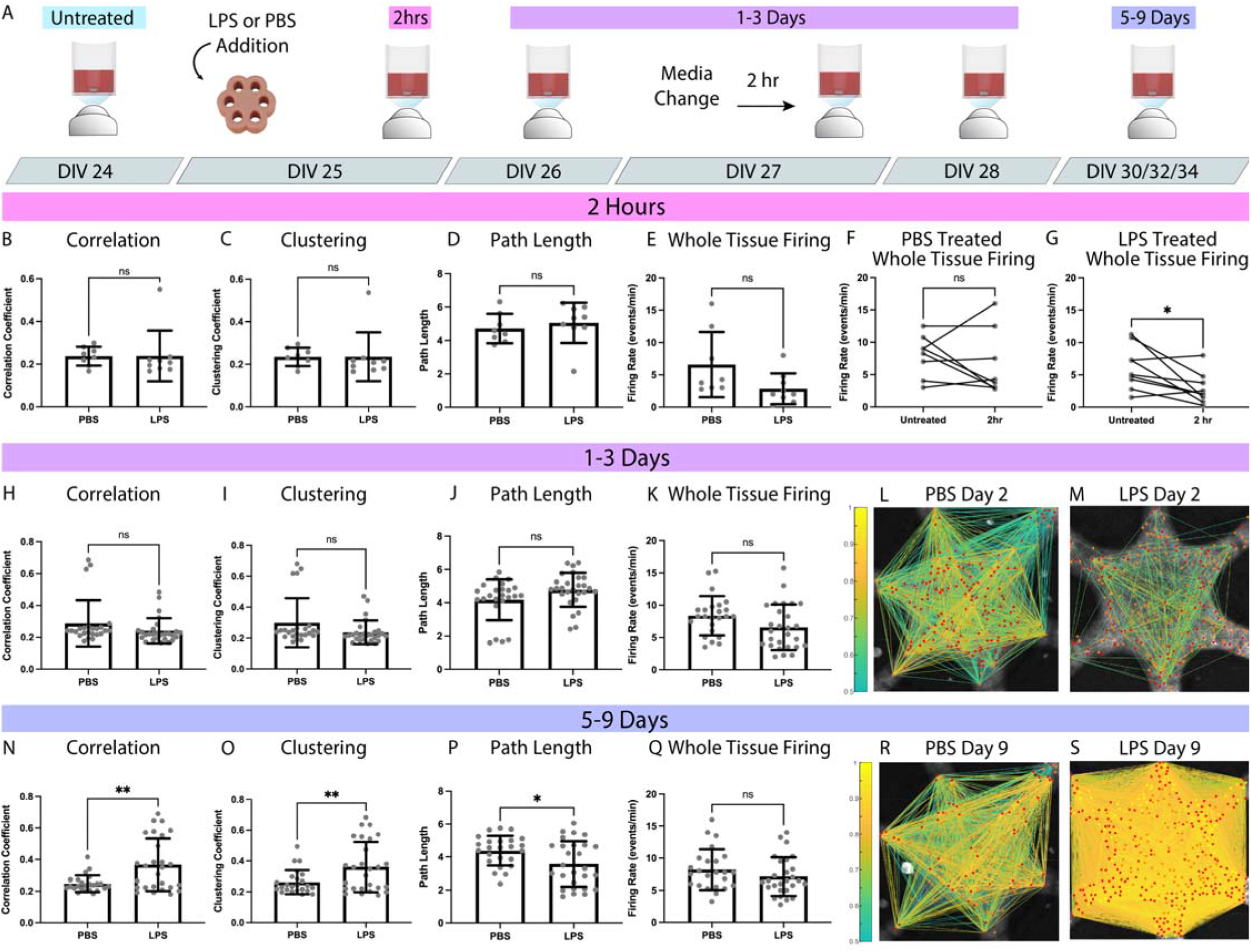
Lipopolysaccharide significantly increases functional connectivity 5-9 days after exposure. (A) Experimental timeline: LPS or PBS was added to the media at DIV 25, allowed to recover for 2 hours, and then imaged (2hr). Subsequent image sessions were done at DIV 26, 27, 28 (1-3 Days) and DIV 30,32,34 (5-9 days). A media change was performed on DIV 27, with a 2-hour recovery before imaging. At 2 hours after exposure, there was no significant difference in average correlation coefficients (p=0.9805, B), clustering coefficients (p=0.9924, C), path length (p = 0.5195, D), or whole tissue firing rate (p=0.0636, E) between PBS and LPS exposed tissues at 2 hrs. The 2 hour treated samples were then compared to matched, untreated samples. The firing rate of PBS samples did not change (p = 0.4479, F), while the LPS samples significantly decreased (p=0.0249, G). At 1-3 days after treatment, there was no statistical difference between PBS and LPS samples in average correlation coefficients (p = 0.1539, H), clustering coefficients (p = 0.0769, I), path length (p=0.0627, J), or whole tissue firing rate (p = 0.0580, K). Example correlational connectomes from representative samples at 2 days after PBS (L) or LPS (M) treatment show similar correlation coefficient distributions above 0.5. At 5-9 days after treatment, LPS samples showed a significant increase in average correlation coefficients (p = 0.0014, N) and clustering coefficients (p=0.0096, O), and a significant decrease in path length (p=0.0198, P) compared to PBS samples. LPS exposure produced no statistical difference (p=0.2126) in firing rate (Q) at 5-9 days. An example correlational connectome from a representative microtissue 9 days after PBS exposure (R) shows similar correlation coefficient distributions above 0.5 to its 2-day counterpart, while the LPS connectome (S) reflects the increase in correlation compared to its 2-day counterpart as well as to the day-matched PBS sample. Significance comparing LPS to PBS samples was determined with unpaired, two tailed t-tests with p<0.05 (^*^p<0.05, ^**^p<0.01). Significance comparing firing rate before and after treatment (F, G) was determined with a paired, two tailed t-test with p<0.05 (^*^p<0.05).

We found no significant changes in correlation (p=0.9805; Figure 5B), clustering (p=0.9924; Figure 5C), path length (p=0.5195; Figure 5D) or whole-tissue firing rate (p=0.0636; Figure 5E) between PBS and LPS treated samples at 2 hours. However, firing rates of individual samples before and after treatment showed no change in PBS firing rate (p = 0.4479; Figure 5F), but a significant reduction in LPS firing rate (p=0.0249; Figure 5G). At 1-3 days, we found no significant differences in LPS and PBS correlation (p=0.1539; Figure 5H), clustering (p=0.0769; Figure 5I), path length (p=0.0627; Figure 5J), or firing rate (p=0.0580; Figure 5K). Correlational connectomes from representative PBS and LPS treated microtissues show connectivity at 2 days post-treatment (Figure 5L, M). At 5-9 days, however, we found significant differences in FC between LPS and PBS treated samples. LPS microtissues showed increased correlation (p=0.0014; Figure 5N) and clustering coefficients (p=0.0096; Figure 5O), and decreased path length (p=0.0198; Figure 5P). These changes in LPS FC were not associated with any changes in firing rate (p=0.2126; Figure 5Q). Correlational connectomes at 9 days show the drastic increase in FC of LPS treated samples over PBS controls (Figure 5R, S).

### LPS interferes with community structure development through disruption of selective functional remodeling

Changes in microcircuit community structure is an indicator of neural dysfunction associated with memory deficits^26,28^. Here, we investigated the influence of LPS-induced acute neuroinflammation on modularity and the underlying community structure of microcircuits in vitro.

At 2 hours after treatment, LPS had no significant effect on modularity (p=0.3074; Figure 6A), intramodular correlations (p=0.3056; Figure 6B), or inter-modular correlations (p=0.056; Figure 6C). Schematic representations of module connectivity (Figure 6D, E; Supplemental Figure 4), show that at 2-hours (DIV25), microtissues have established separation between intra-modular and inter-modular connections in both LPS (p<0.0001) and PBS (p<0.0001) samples. At 1-3 days, modularity of LPS remained the same while PBS showed an upward trend (p=0.0505; Figure 6F). PBS intra-modular correlations showed a significant increase over LPS samples (p=0.0070; Figure 6G), while inter-modular correlations were static (p=0.2690; Figure 6H), suggesting that PBS controls continued to strengthen community structure while LPS samples did not. Schematics of connectivity data (Supplemental Figure 4) show slightly increased intra-modular connections in PBS (intra, p=0.0622; inter, p=0.8413; Figure 6I), while LPS-treated networks remained the same (p=0.5200; p=0.8524; Figure J). At 5-9 days, the overall FC increase in LPS samples was associated with a significant reduction in modularity compared to PBS controls (p<0.0001; Figure 6K). While we found a significant increase in LPS intra-modular correlation (p=0.0133; Figure 6L), there was a disproportionate increase in LPS inter-modular correlation (p<0.0001; Figure 6L, M), effectively reducing network segregation. Schematics of connectivity data show PBS controls maintained well-defined modules at 5-9 days (Figure 6N), while LPS treated samples reduced the strength and separation of modules (Figure 6O). The resulting effect of LPS treatment was the disruption of community structure in microcircuits beginning at 1-3 days, and more robustly at 5-9 days after treatment.

**Figure 6.**
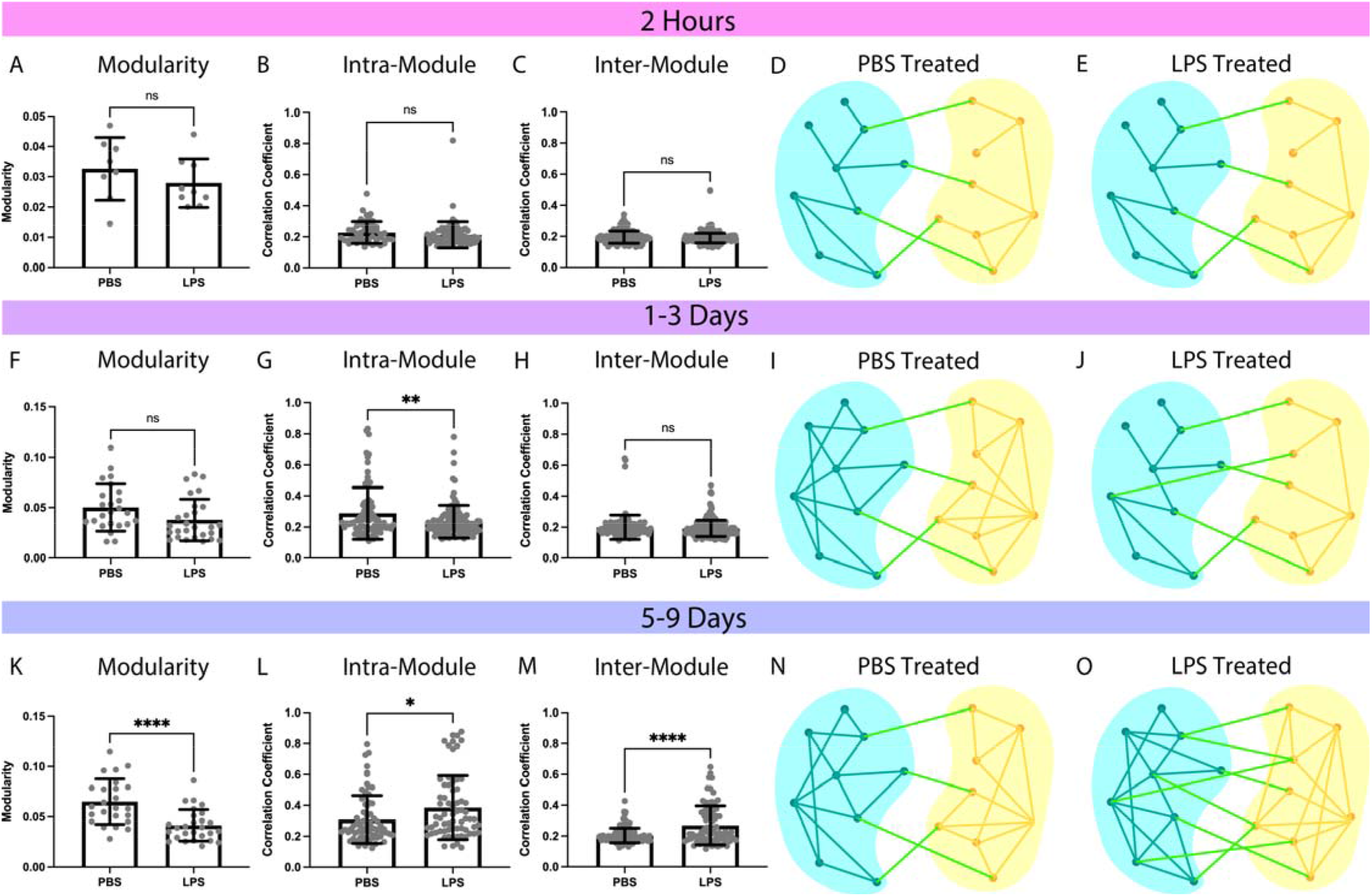
LPS disrupts community structure at 5-9 days after exposure. At 2 hours after treatment, there was no significant difference (p=0.3074) between PBS and LPS in modularity (A), including no difference in the intra-modular (p=0.3056, B) or inter-modular (p=0.056, C) correlation coefficients. Schematic representations of connectivity data (Supplemental Figure 4) display the stronger intra-modular connections in blue (module 1) and yellow (module 2) than inter-modular connection (green) for both PBS treated (p<0.0001, D) and LPS treated (p<0.0001, E) samples at 2 hours, a. At 1-3 days after treatment, there was no significant difference (p=0.0505) between PBS and LPS samples in modularity (F), however, there was a significant increase (p = 0.0070) in intra-modular correlation of PBS (G). Inter-modular correlations remained the same at this time point (p=0.2690, H). Schematics representing of connectivity data (Supplemental Figure 4) shows PBS treated samples (I) with slightly increased in intra-modular connections (p=0.0622) and unchanged inter-modular connections (p=0.8413), while LPS treated samples (J) show no changes from 2 hours (intra, p=0.5200; inter, p=0.8524). At 5-9 days the modularity of LPS samples were significantly lower than PBS samples (p<0.0001, K). There was a significant increase in LPS intra-modular correlations (p=0.0133, L) and a super significant increase in inter-modular correlations (p< 0.0001, M). Schematic representations of modular connectivity data (Supplemental Figure 4). PBS treated samples (N) were unchanged from 1-3 days (p=0.6338), and significantly high intra modular correlations compared to 2 hours (p=0.0082), while LPS treated samples (O) show a large increase in connectivity, which increases inter-modular connections (p<0.0001) and intra-modular connections (p<0.0001). Significance was determined with unpaired, two tailed t-tests with p<0.05 (^*^p<.05, ^**^p<0.01, ^***^p<0.001, ^****^p<0.0001).

## Discussion

Coordinated neural network activity is paramount to a healthy functioning brain and an important feature to examine in vitro. Numerous studies have demonstrated the utility of functional characterization in capturing underlying neural activity during development and disease both in vivo and in vitro ^3,7,15,19–23,25–30^. Because neural cell behavior emerges from the development of complex interconnected networks, extending the functional analysis to include network-wide activity is essential to understanding functional dynamics in vitro. This can be achieved through the application of GTA, which has been used in vivo and in vitro to examine changes in microcircuits to understand early disease states. Here, we demonstrated the application of longitudinal functional imaging and GTA analysis of network dynamics of microcircuits in a 3D primary cortical microtissues. Briefly, we discovered that microtissues undergo significant functional remodeling over weeks of maturation, resulting in the development of community structure which strengthens over time. LPS-induced neuroinflammation significantly increased overall FC of microtissues, which disrupts the existing community structure. Cumulatively, these results have broad implications for the examination of microcircuit dynamics in 3D in vitro models of development and disease.

We assessed network activity of microtissues imaged from DIV14-34, which showed significant functional remodeling marked by weekly fluctuations in FC, measured by correlation, clustering, and path length. Remodeling of FC, while influenced by changes in the underlying physical network structure^19,31^, reflects fast cellular dynamics related to synaptic regulation, including changes to neurotransmitter release, receptor density, and even glial-mediated pruning^55^. This functional remodeling of in vitro microcircuits has significant implications for the complexity of neural networks that arises in the absence of external stimuli^45^. Cultured microtissues constructed strong community structure over weeks of maturation through the formation of highly connected modules. The large changes in FC coincided with selective remodeling of specific node-pair connections, which resulted in higher intra-modular correlations compared to inter-modular correlations. The formation of such modular structure is important in developing specialized neural ensembles involved in basic learning and memory^23,26,46^. Features of FC can assist in inferring information about the underlying structural network^19,31,32^. Structural topology of a network can range from extremely ordered regular networks, to disordered random networks. In the current study, the limited correlation between node distance and correlation coefficient suggests that microtissues contain topology closer to random networks than regular networks. However, the formation of strong community structure indicates that that the network is not random, but more likely scale-free. The presence of a scale free networks would support the formation of an in vivo-like topology associated with cortical and hippocampal development, as opposed to a regular network more associated with cerebellar development^32^. This would differ from 2D cultures, which primarily exhibit features of small world networks skewing toward local connectivity^19,45^. Such small world structure may be a result of the dimensional limitations of neurite growth, and thus connectivity, in 2D models^5^. Further investigation of other topologies, such as the formation of hub nodes and rich-clubs^56^, will give a more complete picture of the structural networks in microtissues during maturation.

While the presence of large scale synchronous bursts are associated with neuronal fate during development, the exact role and progression of the burst rate through maturation has yet to be elucidated^7,20,21,29^. We found that the rate of whole-tissue synchronous bursting was not a sensitive metric of underlying network changes, however we observed changes in firing patterns in individual microtissues over time. This finding is similar to synchronous bursting in 2D cultures, which exhibit hugely variable bursting patterns over time, as well as 3D human neural organoids, which develop increasingly complex, nested bursting patterns during maturation^20,21,57^. These changes in burst patterns likely indicate the development of a flexible interplay between excitatory and inhibitory network inputs, which are associated with underlying mechanisms of attentional neural dynamics and learning^7,57,58^.

The addition of LPS to neural tissue drives acute neuroinflammation by triggering the endotoxin response system, which induces morphological and functional changes in glia^37,59^. Both microglia and astrocyte behavior play a role in synaptic regulation during both normal development^55,60^ and neuroinflammatory disease states, including Alzheimer’s Disease, traumatic brain injuries, and instability of implanted neural devices^22,61,62^. In the current study, we addressed the downstream effects of LPS-induced neuroinflammation on the activity of neurons in a heterogeneous, multicellular, glia-rich environment. Functional dysregulation after LPS exposure was consistent with results from 2D cultures, awake behaving rats, and humans^30,49,51^. A study using 2D primary cultures found diverging effects of LPS on glial-mediated synapse density, where LPS-exposed microglia reduced the number of synapses and LPS-exposed astrocytes increased the number of synapses^63^. This would suggest that increases in FC in our microtissues may be attributed to glial-mediated changes to the structural neuronal network. A benefit of an engineered culture environment is the ability to add or subtract specific cell types. As such, further investigation of the role of different glial cells in these functional changes could improve our understanding of multicellular dynamics associated with neuroinflammatory mechanisms. Further, the disruption of community structure of microcircuits may have broad implications for the role of glia in cognitive impairments exhibited after LPS exposure in both rats and humans^49,50,53,64^.

While the current study demonstrates changes in FC over time and after LPS exposure, we expect to see a more complete view of neural function and dysfunction with longer experimental timelines. Extending the experimental window over months would allow for the examination of age dependent responses to neuroinflammatory drivers and exploration of the limits of community structure formation in mature microtissues. While we demonstrate the use of this platform for functional imaging of neurons, this culture method is compatible with high resolution, longitudinal imaging of morphology and migration of multiple cell types simultaneously within a single sample.

Further, the application of such longitudinal live imaging and functional analysis to primary cultures sourced from transgenic disease models offers potential for capturing early functional changes in specific disease pathologies.

A major pitfall of in vitro culturing platforms is the absence of vasculature^14^. As vascular instability is likely involved in chronic neuroinflammation and neurodegenerative disease^65^, the lack of vasculature in vitro is a significant concern. However, by its nature, an avascular culture model of the brain decouples the systemic and local “innate” immune systems, providing a unique opportunity to study the complex multicellular dynamics of innate neuroinflammation in an experimentally feasible paradigm.

The results of this study demonstrate the utility of a powerful combination of experimental and analytical methodologies to investigate basic cell behavior at the microcircuit level. We revealed progressive selective remodeling over three weeks of imaging, indicating the formation of community structure without external synaptic inputs. We showed that the examination of microcircuits with the use of a GECI is particularly well-suited for 3D cultures and is a robust method of assessing changes to neural behavior and the underlying neurocircuitry in neuroinflammation. The adaptability of the model to any number of live fluorescent probes and cell types, including primary cells from transgenic disease models, makes this methodology especially versatile. The combination of multimodal experimental and analytical tools demonstrated in this work expands the impact of in vitro models in examining single cell and network dynamics underlying neural development and disease.

## Materials and Methods

### Agarose Gels

Agarose microwells were formed in 24-well glass bottom tissue culture plates (P241.5HN, CellVis) with a custom machined injection mold. A 2% (w/v) agarose (16500, Invitrogen) solution in phosphate buffered saline was warmed in a microwave in 30 second increments until agarose powder was dissolved. Hot agarose was drawn into a syringe and pushed through the injection mold into the well and cooled on ice until solid. Solidified agarose inside the injection mold was separated from the agarose outside of the mold before carefully removing the mold from the well. Complete cortical media (CCM) was added over the agarose and incubated at 37C. Media (CCM) was changed on each plate twice, with a minimum 2hr incubation time for each change, before seeding cells.

### Animals

All animals were handled in accordance all animal handling and procedures were carried out in accordance with approved Brown University and Providence VA Medical Center Institutional Animal Care and Use Committee (IACUC) protocols. Timed-pregnant Sprague Dawley Rats were purchased from Charles River Laboratories and monitored for several days before pup litters were birthed.

### Three-dimensional Primary Cortical Cultures

Three-dimensional cortical cultures were formed using primary tissue collected from rats at postnatal day 1. Litters were sexed and an equal number of male and female pups were selected for collection of cortices. Cortices were dissociated following cell isolation protocol adapted from Brainbits. Briefly, animals were euthanized by hypothermia followed by decapitation before whole brain tissue was removed and placed in cold Hibernate A medium (HA, Brainbits) with B27 supplement (17504-044 Invitrogen). Following removal of the dura, the cortex was transected away and stored on ice in Hibernate A + B27 (HA-B27) until dissection of all cortices was complete. Cortices were cut into smaller pieces and added the dissociation solution made from 3mg papain/mL in hibernate solution (PAP, Brainbits) (HACA, Brainbits). Tissues were incubated in dissociation solution at 30C for 30 min with gentle inversion every 5 minutes. The supernatant of papain was then carefully aspirated and replaced with warm HA-B27 followed by trituration with a P1000 pipette tip 20 times. The resulting cell suspension was then dripped through a 40uM filter and washed with additional HA-B27. The filtered solution was centrifuged at 150 rpm for 5 min before careful aspiration of supernatant followed by resuspension of the cell pellet in complete cortical media (CCM). The cell suspension was filtered, spun down and resuspended in CCM two more times. A small aliquot from the final suspension was taken and diluted 1:4 in CCM and then 1:1 in Trypan Blue (T8154, Sigma-Aldrich). A live/dead cell count was determined by hemocytometer and a cell viability was assessed. Only cell batches with >90% viability were used for experiments. Cells were seeded into agarose-molded cell culture plates at 2.7×10^8^ cells/well suspended in 10µl CCM. Any bubbles were removed with a sterile needle and then plates were placed in the incubator for 20 minutes while cells settled. After 20 minutes, 600uL of CCM was carefully added dropwise to each well, avoiding disruption of cells. At 24 hours post seeding, successful self-assembled microtissue formation was verified by the presence of tissue contraction. A 500uL media change of CCM was completed every other day following seeding.

### AAV Transduction

Genetically encoded calcium indicators (GECIs) were introduced into 3D primary microtissues at DIV1 via adeno-associated infection. The GCaMP6s reporter was expressed in neurons under the human synapsin promoter following successful AAV transduction (AAV1-hSyn1-mRuby2-GSG-P2A-GCaMP6s-WPRE-pA, AddGene). Briefly, microtissues were infected at DIV1 by diluting stock viral solution in CCM to a final concentration of 1e6vg/ml and then replacing 1ml of CCM within tissue culture plate well with the virus-containing media. Plates were replaced at 37C, 5%CO2 and left to incubate with viral-containing media for 3 days before performing a complete media change with non-viral CCM.

### Lipopolysaccharide Challenge

Microtissues were exposed to a lipopolysaccharide (LPS) challenge on DIV 25. LPS (L4391, Sigma-Aldrich) was diluted to 10ug/mL in CCM and delivered to microtissues with a full media change. Microtissues were exposed to LPS for 2 days before replacing media with CCM on DIV 27.

### Calcium Imaging

Microtissues were imaged within the glass-bottom plate on a confocal laser scanning microscope (Olympus FV3000) in 4-min recording sessions. Briefly, the glass-bottom plate was placed on the microscope stage with the multi-well plate adapter. With the microscope aperture set to 800um (fully dilated), the 488nm laser and EGFP filter were used to visualize fluorescent expression and bring the microtissue within the focal plane of the 10x objective. The 4-min time-lapse videos were recorded at 15 frames per second. Plates were imaged for no more than 30 minutes at a time before being placed back in the 37C, 5%CO2 incubator. Microtissues were fed every other day and imaged on non-feeding days between DIV 14-34. For LPS or PBS treated samples, microtissues that were imaged on feeding days (DIV 25 and 27) were given 2 hours to recover in the incubator post-feeding.

### Microtissue Fixation and Removal from Agarose

Microtissues were fixed within the agarose-molded wells of the tissue culture plate with cold PFA solution (4%(w/v) paraformaldehyde, 8% (w/v) sucrose) (F79-500, Sigma-Aldrich) (S7903, Sigma-Aldrich). Microtissues were fixed for 2 hours followed by 3 PBS washes. Following several PBS washes, the agarose mold was carefully removed from the bottom of the plate well and the excess gel was cut away while remaining cautious to not cut too close to the tissue itself. The remaining agarose mold containing the microtissue was then placed in warmed PB and heated on a hot plate at 200C for 10 minutes with frequent agitation until the agarose had melted, leaving behind the freely floating fixed microtissue. The microtissue was then transferred to a 35mm petri dish and washed several times with warmed PBS to remove any residual agarose before transfer to a 48-well plate. Microtissues were left in cold PBS until immunohistochemical labeling was performed.

### Immunohistochemistry

Following fixation and agarose removal, the microtissues were immunoassayed for cell-specific antigens. 300 µl of block solution (10 mL solution consisting of 1 mL normal goat serum, 8.9 mL 1xPBS, 0.1mL TritonX-100, and 0.4 g of Bovine serum albumin) (005-000-121, Jackson Immuno Research) (T8787, Sigma-Aldrich) (A7030, Sigma-Aldrich) was added to each 48-well containing a microtissue and placed on a shaker with gentle agitation for 2 hours. Following the 2-hour block, solution was removed and 300 µl of a primary antibody cocktail diluted in blocking solution was added to each well and placed back on the shaker for room temperature incubation overnight. The primary antibody cocktail consisted of neuronal nuclei marker NeuN (mouse anti-NeuN, MAB377, 1:500, Milipore), glial fibrillary acidic protein (GFAP) (Chicken anti-GFAP, AB5541, 1:200, Milipore), and microglial marker, ionized calcium-binding adapter molecule-1 (Rabbit anti-Iba, 019-19741, 1:200, WAKO Chemicals). Following the overnight incubation in primary cocktail, the antibody solution was removed and microtissues were subsequently washed two times for 2 hours with PBT (50 mL solution consisting of 49.9 mL of 1x PBS, 0.1mL of Triton X-100). Then, 300 µl of blocking solution was again added to each well and left to shake for 2 hrs. Following block, 300 µl of a secondary antibody cocktail diluted in blocking solution was added to the wells and left to shake at room temperature overnight. The secondary antibody cocktail included goat anti-mouse CY3 (115-165-068, 1:800, Jackson Immuno Research), goat anti-chicken Alexa Fluor 488 (A11039, 1:800, Invitrogen), and goat anti-rabbit Alexa Fluor 635 (A31577, 1:800, Invitrogen). On the third day, the secondary antibody cocktail solution was removed and microtissues were washed with PBT four times over four hours. Following the PBT washes, the dsDNA fluorescent marker DAPI (D1306, 1:1000, Invitrogen) was diluted in PBT and added to microtissues and left to incubate for one hour.

DAPI solution was removed, and each well was replaced with 500 µl of PBS. Microtissues were transferred to 35mm glass bottom petri dishes for confocal imaging or placed at 4C for storage.

### Video preprocessing

Calcium videos (.oir format) were loaded into FIJI using the Olympus Viewer plugin (https://imagej.net/OlympusImageJPlugin). Video files were first saved as tiff stacks for later processing. Maximum projections across the time domain were created with the “Z Projection” function, with the “Max Projection” option selected to identify any cell bodies expressing calcium fluorescence during the video. Max projections were then saved as .tiff files for semi-automated ROI detection.

### Semi-Automated ROI Detection

ROI detection was performed with Python code (ca_roi_analysis: https://github.com/neuromotion/calcium-roi-analysis). The max projection tiff image was loaded by adding it to the path in the current.yaml file. Images were smoothed by a Gaussian filter with a kernel of a sigma that corresponded to the size of the cell bodies (sigma = 2.5). Smoothed images underwent adaptive thresholding based on the intensity of local neighborhoods (neighborhood size, threshold_block_size=5) using a curated threshold fraction above the mean intensity (threshold, threshold_fraction_above_mean=0.008-0.018) to create a binarized image. The binarized image was then used to make a distance transform image. A Laplacian of Gaussian pyramid was applied to the distance transform image with a range of scales (min_sigma=1, max_sigma=5) to detect “blobs”, or cells, of varying sizes with a local threshold (local intensity threshold, blobs_log_thresh =0.01). Manual interactive correction of ROIs was achieved using the napari visualizer, and any ROIs of unattached cells outside of the microtissue boarder were removed. ROI seed positions were saved as an .npz file.

### Calcium Signal Extraction

Calcium signals were extracted from ROI’s by first making a mask of ROI locations (CalciumMask: https://github.com/neuromotion/CalciumMask). ROI seed positions were imported from the Python code output (.npz file) and overlaid onto the max projection image. Then a 3-pixel radius circle ROI was place over each seed position and a mask of all ROIs was created. This mask was then placed over each frame of the calcium video and exported as a tiff stack. Calcium signals were extracted from masked videos using the FluoroSNNAP^40^ application (https://www.seas.upenn.edu/~molneuro/software.html). Masked tiff files were segmented with the “Segmentation GUI” using the threshold function to find ROIs from the masked video. In analysis preferences the analysis modules were set to “convert raw fluorescence” and “detect calcium transients” in order to extract single cell calcium traces and events. Acquisition frame rate was set to the image sampling rate of 15Hz. Df/f traces were created by detecting the baseline fluorescent signal, calibrated from the 20^th^ percentile of the signal across a 10 second time window. Calcium events of individual ROIs and whole tissues were detected with a template-based method, which identifies events by calculating similarity between a moving window of the calcium trace to library of calcium waveform templates. Events were detected with a similarity threshold of 0.7 and a minimum amplitude of 0.01. Calcium transients were extracted in batches and outputs were saved in the processed_analysis.mat file.

### Graph Theory Analysis

MATLAB code was used to create a Pearson cross-correlation matrix from the single-cell calcium transients extracted from the FluoroSNNAP output (CalciumSignalProcessing:https://github.com/neuromotion/CalciumSignalProcessing). Pearson cross-correlation was calculated using the MATLAB corr() function, and average correlation was calculated by the mean cross-correlation value. Whole-tissue firing rate was calculated by the number of whole-tissue events (detected by FluoroSNNAP) divided by the 4-min recording time. The cross-correlation matrix was used as the input for the network-analysis-master code from Dingle et al.^7^ (https://github.com/yutingdingle/network-analysis), which compiled previously published code to calculate the average clustering coefficient, average pathlength, average modularity, and identify the number of modules and node compositions of each module. In this code, clustering coefficients were calculated using a threshold-free method from Onnela et al.^43^ which used weighted graphs to calculate intensity of local network connections, limiting the pseudo correlation effect of thresholded graphs. The pathlength of the weighted network was determined by the shortest distance between nodes (1/correlation coefficient) proportional to the size of the network (1/N(N-1), where N is number nodes in the network) from Muldoon et al^44^. Module detection was achieved by hierarchical clustering of the cross correlations and used to calculate the modularity (sum of the correlation coefficients in a module / sum of the expected correlation coefficients in a random network) from M.E.J. Newman^47^.

### Contraction Analysis

Contraction analysis was performed using custom MATLAB code (MicrotissueContraction https://github.com/neuromotion/MicrotissueContraction). First, the pixel diameters of the agarose pegs were measured with the PegMeasurment.m code by hand curating peg edges from at least three different phase images. The average peg diameter from these measurements was used to automatically identify peg positions in the Contraction.m code. In this code, image file paths were set to the “Path” variable, images were binarized using the imbinarize() function, and peg positions were identified using the edge() function with Canny edge detection. Peg centroids were then used as static landmarks in the image to place an ROI over the center of the microtissue and a mask of the center ROI was created. Due to variability in image contrast, some images produced poor binarization of the tissue edge. This was solved by a combination of thresholding and hand curation of the binarized tissue edge. Binarized images, which showed tissue in white and background in black, were then used to calculate tissue contraction away from the pegs was measured by the percent of white (tissue positive) pixels within the ROI.

## Statistical Analysis

Statistical analysis was performed using GraphPad Prism software. All data was collected from microtissues across 3 biological replicate litters. Control matched samples were grouped by week and statistical significance of functional metrics including correlation coefficient, clustering coefficient, path length, modularity, and whole tissue firing rate were tested by one-way ANOVA with Tukey HSD post-hoc analysis, where each data point represented a single recording of a microtissue within the grouped week Statistical significance of functional metrics between PBS and LPS treated samples was tested by unpaired, two-tailed t-test, where each data point represented a single recording of a microtissue with the grouped time frame after exposure. Statistical significance of intra-modular and inter-modular correlation was tested by unpaired, two-tailed t-test, where each data point represented the average correlation of a single module or module-pair. Significance level of alpha<0.05 was determined (^*^p<0.05, ^**^p<0.01, ^***^p<0.001, ^****^p<0.0001). All error bars represent the standard deviation from the mean.

## Data Availability

All data generated or analyzed during this study will be made publicly available upon publication and can be made available upon request.

## Acknowledgments

The authors thank Geoff Williams for his guidance and technical support with image acquisition, Ben Goddard for his guidance and technical support on software development, Diane Hoffman-Kim and Liane Livi for their cell culture guidance and scientific support.

## Author Contributions

E.A., S.B. and D.A.B. conceived of the experiments; E.A. and S.B. performed the experiments; E.A., S.B. and E.P. performed image acquisition and data analysis; E.A. M.I.R., and E.P. wrote the MATLAB and Python code; E.A. wrote the first draft of the manuscript; All authors reviewed and edited the final manuscript.

