## Supplementary figures for "Lipopolysaccharide-induced neuroinflammation disrupts functional connectivity and community structure in primary cortical microtissues"

**Supplemental Figures:**

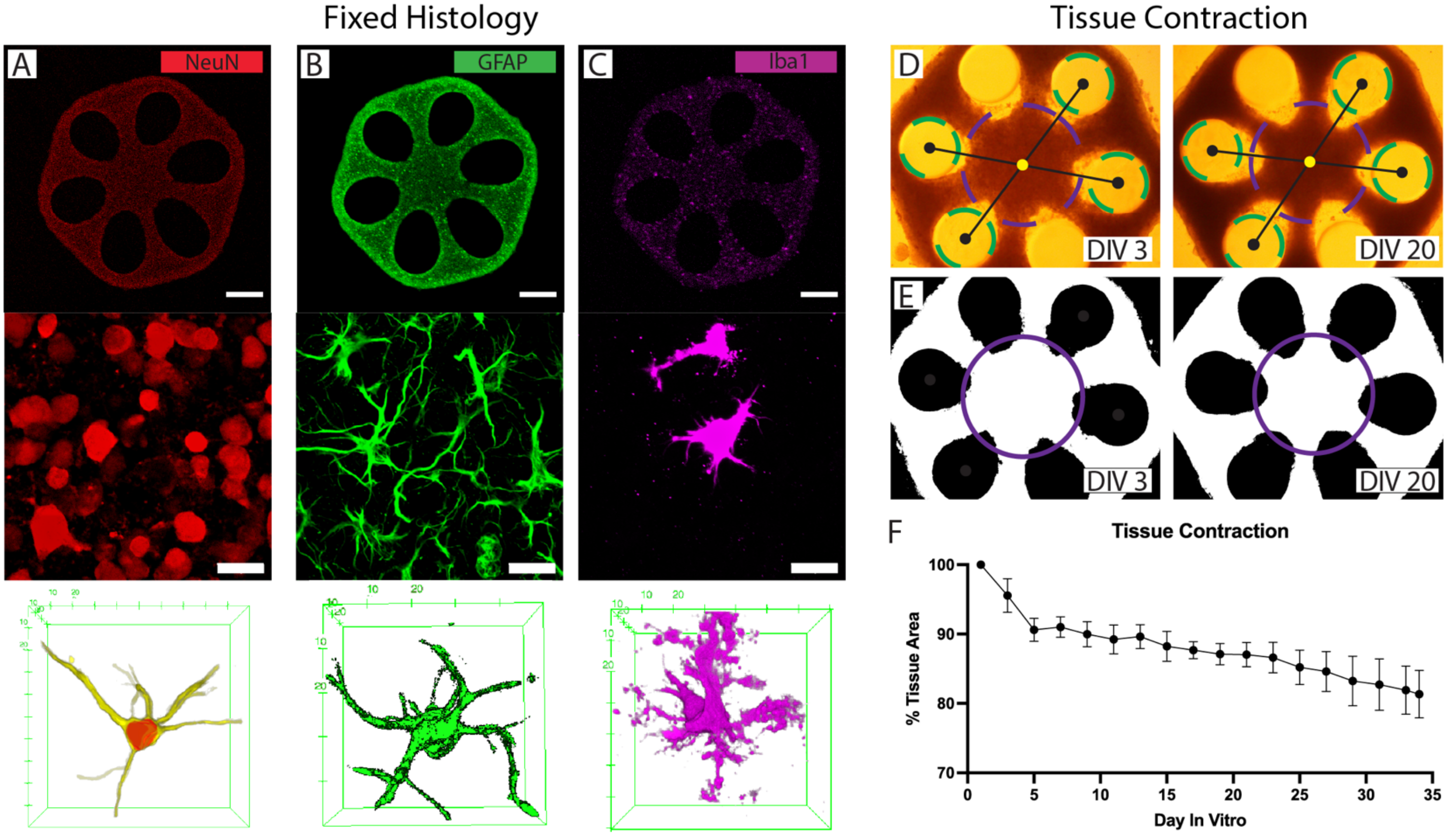


**Figure S1. Microtissues morphology.** Microtissues exhibit 3D morphology of multiple cell types, including (A) neurons (NeuN, red), (B) astrocytes (GFAP, green), and (C) microglia (Iba1, magenta). Microtissues show macromorphology changes characterized by tissue contraction over weeks (D). Contraction of the center of the microtissue was measured with custom MATLAB code by placing an ROI in the center of the tissue (purple), binarizing the image (E) so that the tissue is white, and then measuring the percent of white pixels within the ROI. (F) Microtissues contracted the most within the first 14 days in vitro (89.63% tissue area), and continued to contract through day 34 (81.34% tissue area).

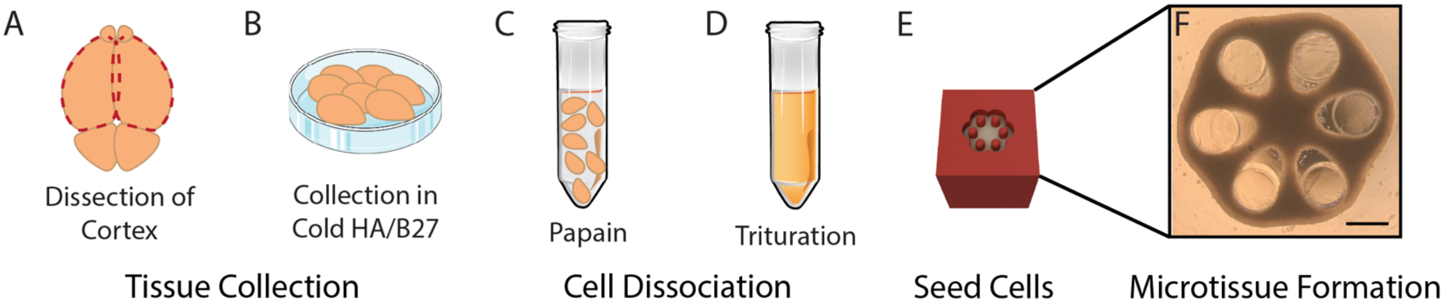


**Figure S2. Tissue collection and dissociation.** (A) Cortical tissue was dissected from P1 rat brains, and (B) placed in cold HA-B27. (C) Cortices were chemically dissociated in papain at 30C for 30 minutes. (D) Physical trituration was used to make a single cell suspension. (E) Cells were seeded into the agarose microwell, (F) forming a microtissue. Scale bar = 500um.

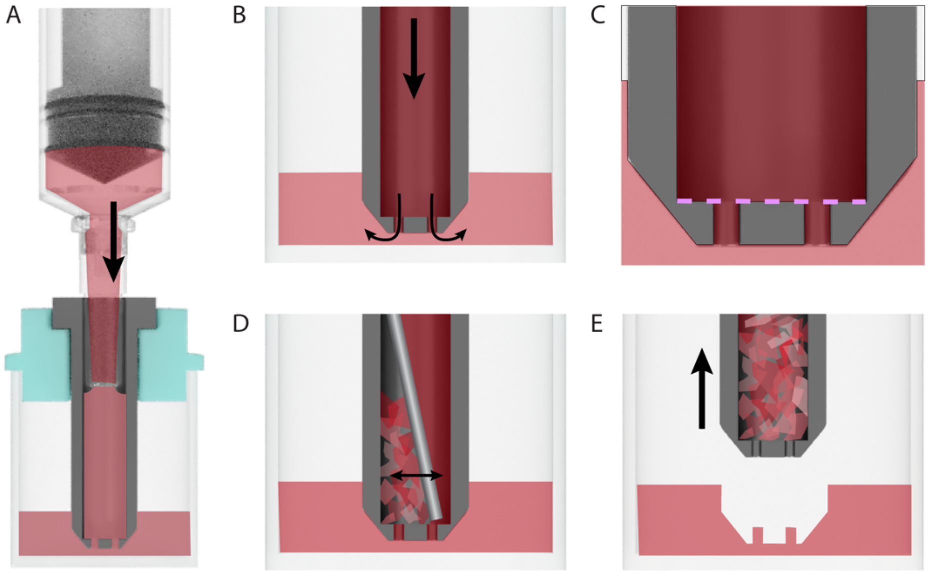


**Figure S3. Agarose microwell formation.** (A) Hot agarose was pushed through the injection mold with a luer lock syringe. (B) Agarose moves from the injection mold into the well through the peg holes. (C) To remove the mold from cooled solid agarose, the pegs need to be separated from the agarose inside the injection column. (D) A sterile metal rod was used to scrape away the agarose inside the injection column, (E) allowing for removal of the mold.

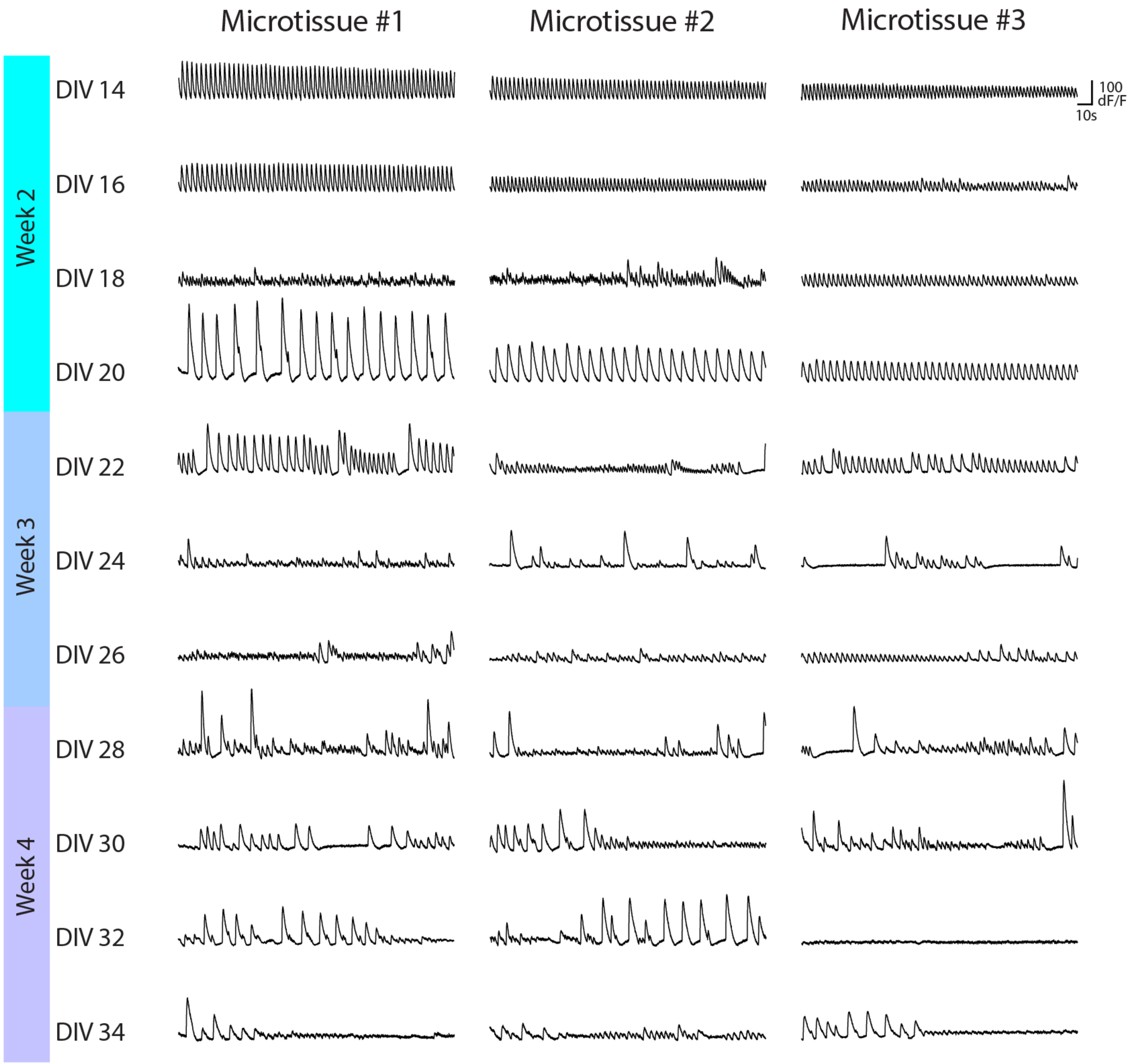


**Figure S4. Calcium traces from three example microtissues.** Each trace is a 4-minute recording from a microtissue. Designated week groupings of recordings are indicated by the row headers of Week 2 (DIV14-20), Week 3 (DIV21-27), Week 4 (DIV28-34).

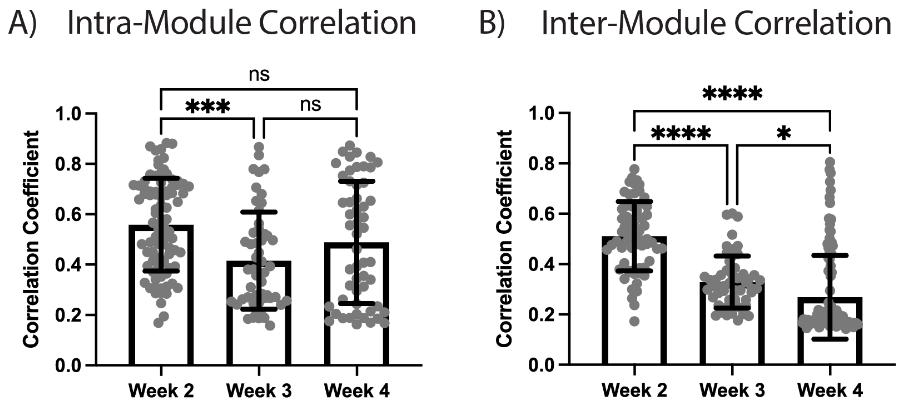


**Figure S5. Progression of module correlations over weeks.** (A) Intra-modular correlations from Week 3 through Week 4. (B) Inter-modular correlations from Week 2 through Week 4.

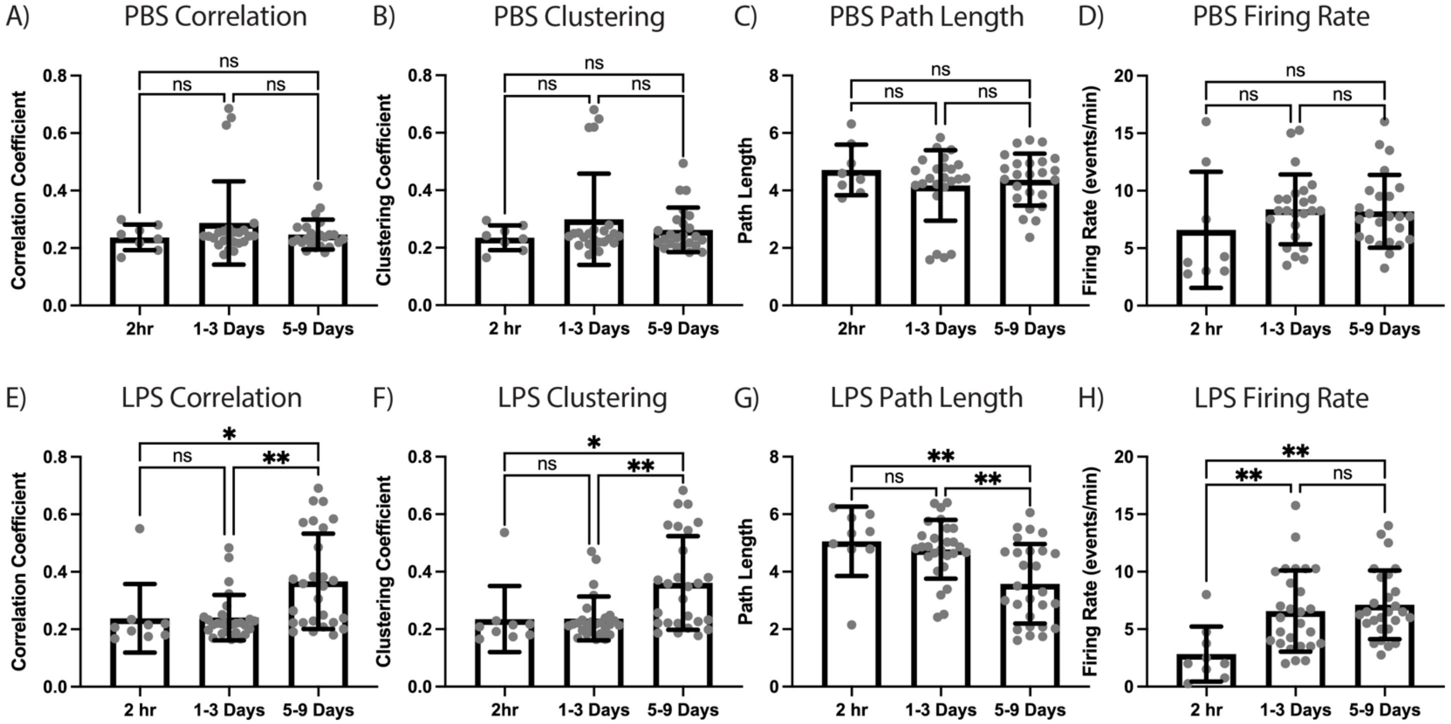


**Figure S6. Progression of module correlations in PBS and LPS treated samples over days.** PBS treated samples show no significant changes from 2 hours to 9 days in (A) correlation, (B) clustering, (C) path length, and (D) firing rate. LPS treated samples exhibit significant changes in (E) correlation, (F) clustering, and (G) path length at 5-9 days, while (H) firing rate was primarily affected at the 2-hour time point.

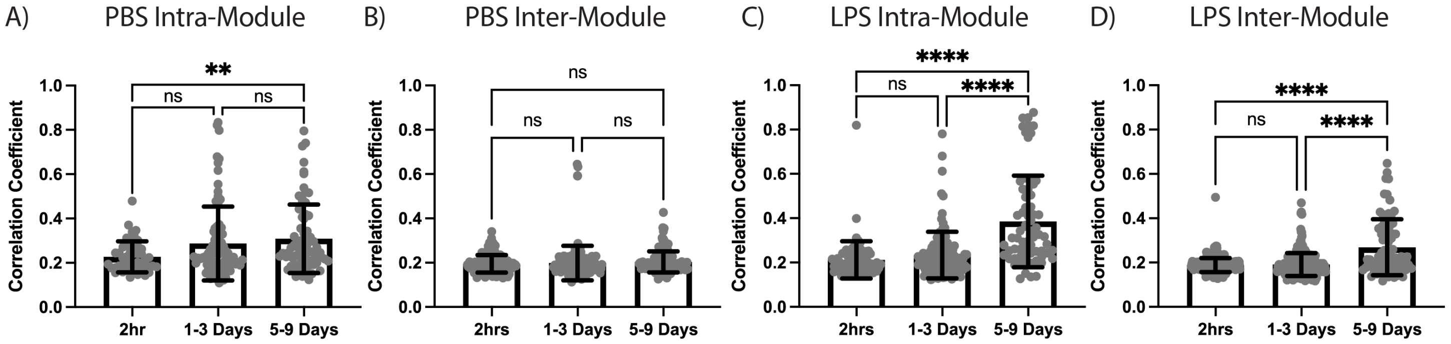


**Figure S7. Progression of module correlations in PBS and LPS treated samples over days.** PBS treated samples show a progressive increase in intramodular correlations from 2 hours to 9 days (A), and no change in inter-modular correlations (B). LPS intra-modular (C) and inter-modular (D) correlations significantly increase at 5-9 days.
